# Rapid selection of P323L in the SARS-CoV-2 polymerase (NSP12) in humans and non-human primate models and confers a large plaque phenotype

**DOI:** 10.1101/2021.12.23.474030

**Authors:** Xiaofeng Dong, Hannah Goldswain, Rebekah Penrice-Randal, Ghada T. Shawli, Tessa Prince, Maia Kavanagh Williamson, Nadine Randle, Benjamin Jones, Francisco J Salguero, Julia A. Tree, Yper Hall, Catherine Hartley, Maximilian Erdmann, James Bazire, Tuksin Jearanaiwitayakul, ISARIC4C investigators, Malcolm G. Semple, Peter J. M. Openshaw, J. Kenneth Baille, Stevan R. Emmett, Paul Digard, David A. Matthews, Lance Turtle, Alistair Darby, Andrew D. Davidson, Miles W. Carroll, Julian A. Hiscox

**Affiliations:** Institute of Infection, Veterinary and Ecological Sciences, University of Liverpool, UK; NIHR Health Protection Unit in Emerging and Zoonotic Infections, Liverpool, UK; School of Cellular and Molecular Medicine, University of Bristol, UK; UK Health Security Agency, Porton Down, UK; Department of Microbiology, Mahidol University, Thailand; Department of Respiratory Medicine, Alder Hey Children’s Hospital, Liverpool, UK; National Heart and Lung Institute, Imperial College London, UK; The Roslin Institute, University of Edinburgh, UK; Royal United Hospitals Bath NHS Foundation Trust, UK; Bristol Medical School University of Bristol, UK; Nuffield Department of Medicine, University of Oxford, UK; A*STAR Infectious Diseases Laboratories (A*STAR ID Labs), Agency for Science, Technology and Research (A*STAR), Singapore

## Abstract

The mutational landscape of SARS-CoV-2 varies at both the dominant viral genome sequence and minor genomic variant population. An early change associated with transmissibility was the D614G substitution in the spike protein. This appeared to be accompanied by a P323L substitution in the viral polymerase (NSP12), but this latter change was not under strong selective pressure. Investigation of P323L/D614G changes in the human population showed rapid emergence during the containment phase and early surge phase of wave 1 in the UK. This rapid substitution was from minor genomic variants to become part of the dominant viral genome sequence. A rapid emergence of 323L but not 614G was observed in a non-human primate model of COVID-19 using a starting virus with P323 and D614 in the dominant genome sequence and 323L and 614G in the minor variant population. In cell culture, a recombinant virus with 323L in NSP12 had a larger plaque size than the same recombinant virus with P323. These data suggest that it may be possible to predict the emergence of a new variant based on tracking the distribution and frequency of minor variant genomes at a population level, rather than just focusing on providing information on the dominant viral genome sequence e.g., consensus level reporting. The ability to predict an emerging variant of SARS-CoV-2 in the global landscape may aid in the evaluation of medical countermeasures and non-pharmaceutical interventions.

## Introduction

There are many distinct lineages of SARS-CoV-2 currently circulating worldwide and some that have become extinct ^1^. Sequence data show that that the genome of SARS-CoV-2 is changing as the pandemic continues. Replication and transcription of the SARS-CoV-2 genome directly drives three types of genetic change in the virus. The first is recombination, and this is a natural consequence of the way in which the virus synthesizes its subgenomic messenger RNAs (sgmRNAs). This may account for insertions and deletions, for example observed in and around the furin cleavage site in the spike glycoprotein ^2^ and other genes ^3^. The second driver of genetic change is the continual accruing of point mutations. These changes may confer advantages in transmission, such as the A23402G, encoding the D614G substitution in the spike protein ^4^, which has come to predominate in global SARS-CoV-2 sequences since the start of the outbreak ^5^. Such point mutations may be driven by errors during RNA synthesis by the viral encoded RNA dependent RNA polymerase (NSP12) and larger replication complex and/or by host mediated processes ^6, 7^. The third mechanism is the potential generation and selection of new transcription regulatory signals (TRSs) and the synthesis of new viral sgmRNAs and proteins ^8^. Promiscuous recombination and mutation in coronaviruses may allow these viruses to overcome selection pressures, transit population bottlenecks and result in the emergence of new variants ^9, 10^.

This variation exists in individual humans/animals infected with SARS-CoV-2, where there will be a dominant viral genome sequence(s) with minor genomic variants ^10^. These latter genomes will have both synonymous (non-coding) and non-synonymous (coding) variations (changes) around the dominant viral genome sequence. These variations may be selected for and become the dominant viral genome sequence when the virus enters a new host, as has been demonstrated with the adaptation of Ebola virus in a guinea pig model of infection ^11^. Alternatively, the variation may exist at a minor variant level but nevertheless impact upon virus biology, for example with the Ebola virus RNA dependent RNA polymerase (L protein) and the relationship with overall viral load in patients with Ebola virus disease ^12^.

Since the start of the COVID-19 pandemic different dominant viral genome sequences and non-synonymous changes appear to rise and fall in the SARS-CoV-2 global sequences ^1^. The D614G spike protein variant of SARS-CoV-2 was first observed in February 2020 and by May 2020 approximately 80% of viruses sequenced contained this substitution. The major clade containing D614G (Pango lineages B.1 and sub-lineages) contained potentially linked substitutions, including C14407U in NSP12 that confers a P323L substitution. However, some lineages, such as A.19 and A.2.4, gained D614G in the spike protein but not P323L in NSP12 ^13^.

Therefore, whether P323L in NSP12 conferred a fitness advantage and was subject to selection pressure is unknown. To investigate the within host selection pressure for the P323L variant, sequential samples from patients with COVID-19 prior to and during the D614G/P323L change in the UK were sequenced to study both the dominant viral genome sequence and minor variant genomes. Additionally, a lineage B SARS-CoV-2 with 323L and 614G in the minor variant population was used to infect two non-human primate models ^13^, cynomolgus (*Macaca fascicularis*) and rhesus (*Macaca mulatta*) macaques. Longitudinal sampling indicated that 323L became part of the dominant viral genome sequence, but not 614G. Reverse genetics analysis of P323L in the background of a 614G virus indicated that the 323L variant grew with a larger plaque phenotype. Overall, this change provided an additive advantage to D614G in the spike protein. In the wider context the work indicated that an emerging dominant sequence could be predicted by analysis of minor variant genomes.

## RESULTS

### Identification of a P323L substitution in NSP12 in the same human patient

To identify whether the P323L substitution occurred rapidly in NSP12, nasopharyngeal swabs were identified in the ISARIC-4C biobank that were obtained from patients infected with lineage B SARS-CoV-2 prior to the major shift from P323 to 323L and D614 to 614G. Samples were further down selected based on clinical information providing a dates of symptom onset, first sample and subsequent longitudinal samples. This provided samples from a total of 472 nasopharyngeal swabs. RNA was isolated from the swabs and used as templates for the amplification of SARS-CoV-2 genome and sgmRNAs using both short (ARTIC-Illumina) and longer-read length (Rapid Sequencing Long Amplicons-Nanopore, RSLA-Nanopore) ^14, 15^. Longitudinal samples from 12 patients had sufficient read depth to call a consensus for the dominant viral genome sequence in each sample and to derive information on the frequency of minor genomic variants, focusing on codon 323 in NSP12 and 614 in the spike protein. In one patient, who was admitted to the intensive care unit at the Royal Liverpool Hospital, both sequencing approaches indicated that the P323L and D614G substitution occurred in the SARS-CoV-2 genome between the 1^st^ sample and 2^nd^ samples taken two days apart (Figure 1A and 1B, respectively). To independently confirm this observation, the source RNA was Sanger sequenced with primers to generate longer amplicons around the potential substitution sites. The data validated that for NSP12 the codon encoding the amino acid at position 323 changed from CCU (encoding P) to CUU (encoding L) (Figure 1C). For the spike protein, the codon encoding the amino acid at position 614 changed from GAU (encoding D) to GGU (encoding G) (Figure 1C). Therefore, the data suggested that both P323L and D614G were rapidly selected in the patient over a two-to-three-day period. Another possibility is that the patient was infecte with a P323/D614 variant and subsequently became infected with a 323L/614G variant through nosocomial infection in the hospital setting. However, we consider this possibility unlikely; a this patient was one of the first cases admitted to the intensive care unit of Liverpool University Hospitals, when there were relatively few other patients present in the hospital at that period of the containment phase.

**Figure 1.**
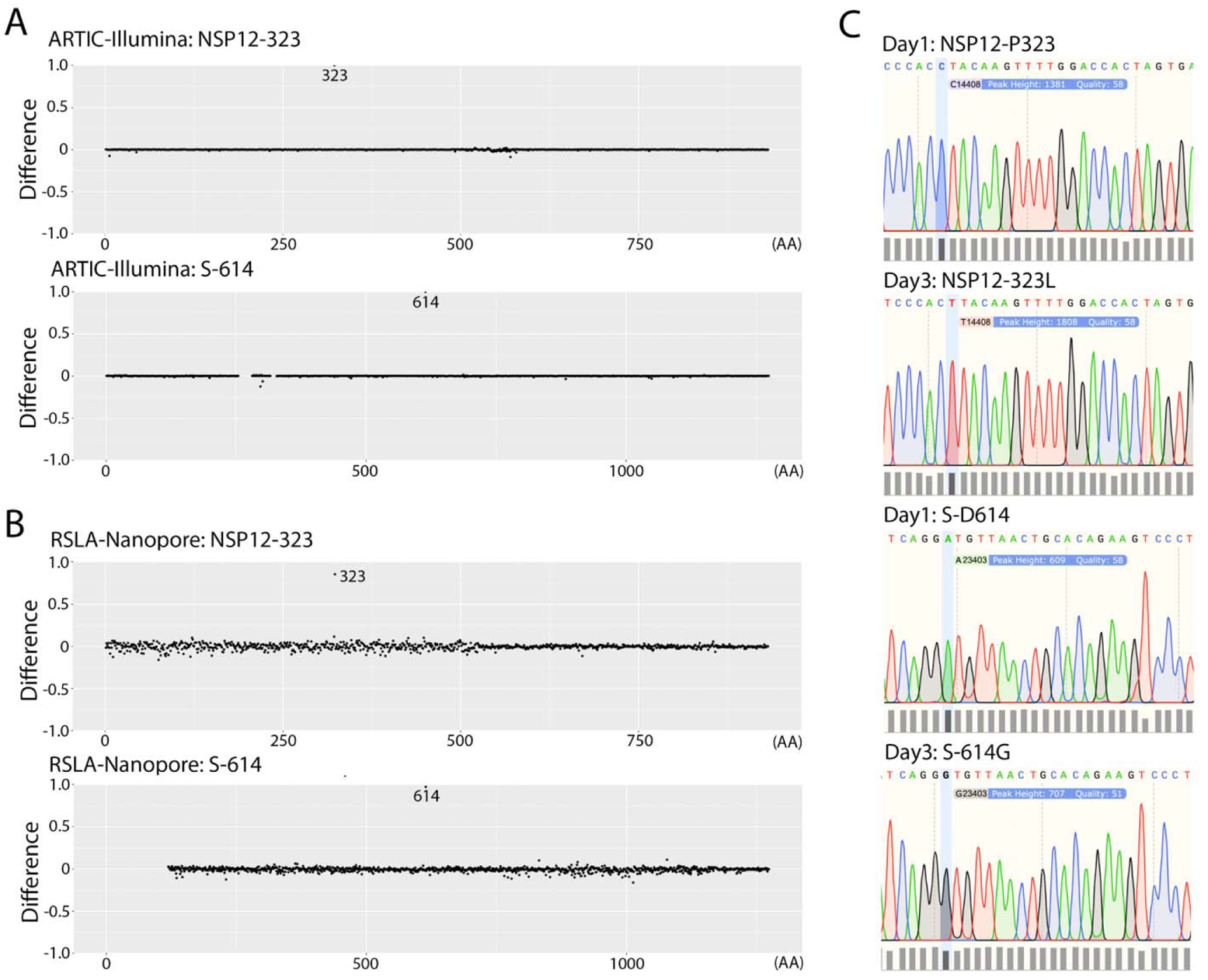
Sequence analysis and amino acid substitution in NSP12 (P323L) and the spike protei (D614G) between an initial sample and one taken two days later in a single patient. Three different sequencing approaches were used: (A) an ARTIC -Illumina approach and (B) an RSLA-Nanopore approach. Individual dots represent a codon position on either NSP12 or the spike protein compared to the Wuhan reference sequence. The difference between the sampling day is indicated by a positive difference indicating divergence of the day 3 sequence away from the Wuhan-Hu-1 complete genome reference sequence (NC_045512) and a negative difference indicating divergence of the first sample taken towards the Wuhan reference sequence. In both cases considering the ratio of a particular position for dominant viral genome sequence versus minor variant. (C) Sanger sequence analysis of the amplicons used to investigate the dominant viral genome sequence around the sites within NSP12 (codon 323) and spike protein (codon 614 that changed between the first and third days of sampling in a patient hospitalized with COVID-19.

The distribution of P323L and D614G at the minor genomic variant level was evaluated in the human population between January 2020 and June 2020, when these substitutions became part of the dominant viral genome sequence. SARS-CoV-2 was sequenced from nasopharyngeal swabs sampled from 522 patients over that time and usable data obtained from 377 (Figure 2).

**Figure 2.**
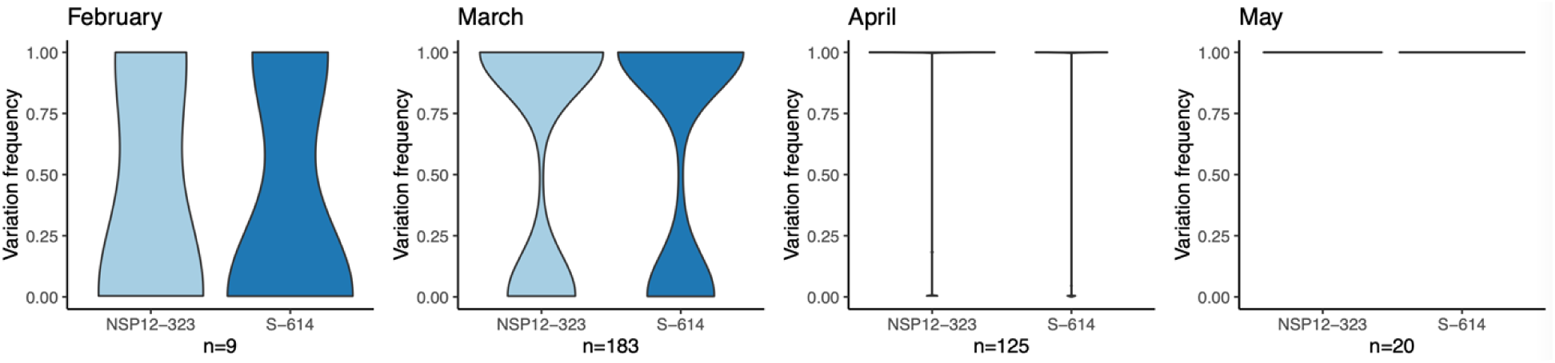
Analysis of the ratio of P323L (light blue) and D614G (blue) at a dominant viral genom sequence and minor variant genomes in 377 patients between February 2020 and May 2020 in the UK. SARS-CoV-2 sequence was obtained from nasopharyngeal swabs from 377 hospitalized patients. The width of the violin plot indicates the number of samples/patients with the frequency on the y-axis. The data shows the transition from P323L and D614G over time in the minor variant genomes, such that by April 2020 in the UK, the 323L and 614G substitutions were part of the dominant viral genome sequence and by May 2020, there was no evidence of P323 and D614.

The data (Figure 2) indicated that there was increasing prevalance from P323 to 323L and D614 to 614G in the February to March sampling period. For both February and March 2020, patients had mixed populations of P323L and D614G. However, for the patients sampled in April and May 2020 the dominant viral genome sequence in each patient had 323L and 614G, suggesting either strong selection pressure and/or multiple founder effects.

### Longitudinal analysis of variation in non-human primates and cell culture

To investigate whether the P323L substitution was driven by strong selection pressure, nasopharyngeal swabs were taken longitudinally from cynomolgus and rhesus macaques (12 animals of each species, a mix of males and females) that had been infected with an isolate of SARS-CoV-2 prior to the P323L and D614G changes; SARS-CoV-2 Victoria/01/202040, that had been sampled on the 24^th^ January 2020 ^16^. The isolate had been passaged three times in cell culture to generate stock virus prior to infection of the cynomolgus and rhesus macaques. Sequencing of the stock virus indicated a very low proportion of NSP12 323L and spike 614G (Figure 3).

**Figure 3.**
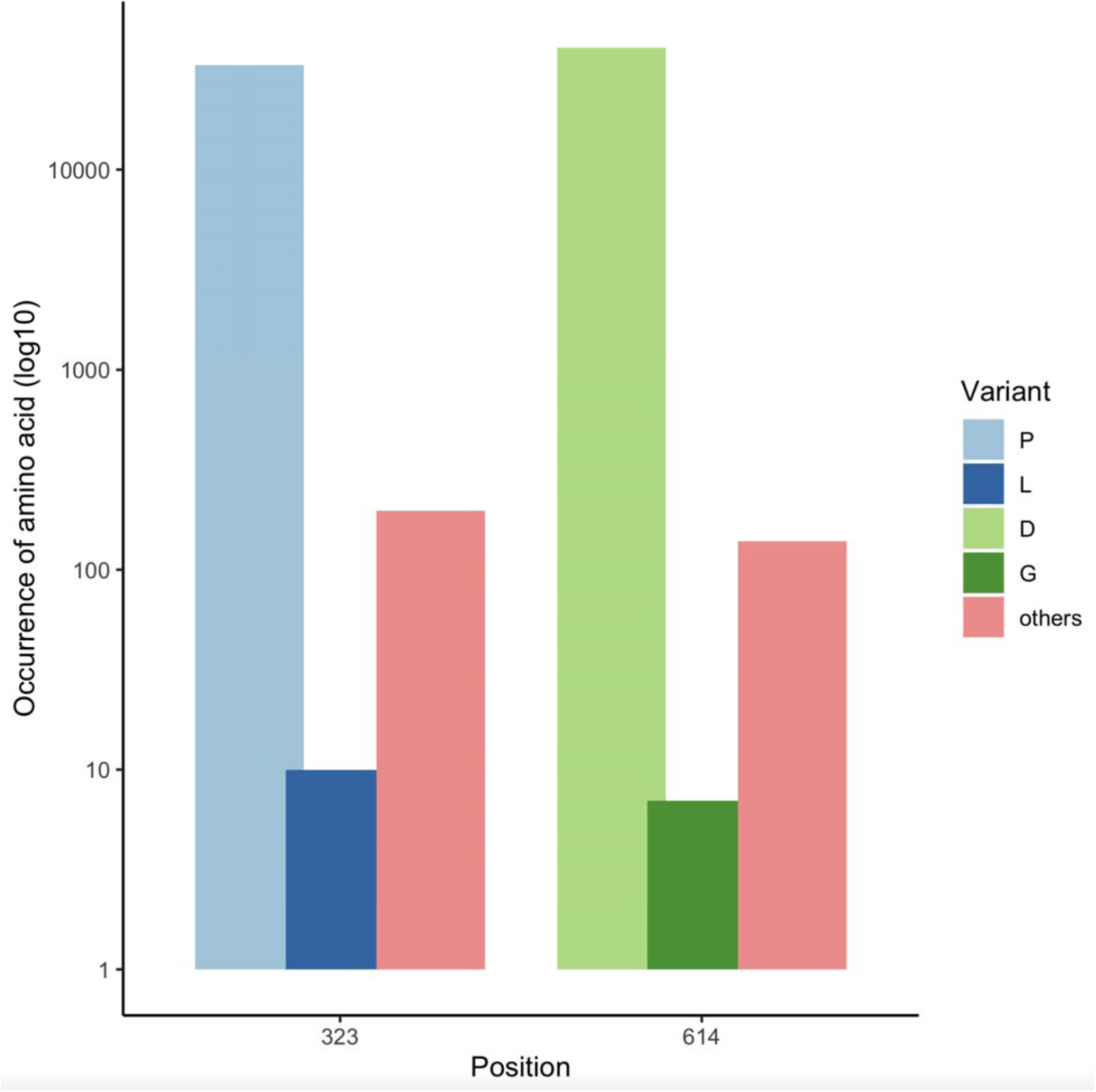
Histogram showing the amino acid coverage at position 323 in NSP12 and 614 in th spike protein in the SARS-CoV-2 Victoria/01/202040 stock as determined by ARTIC-Illumin sequencing. Site coverage is shown on the y-axis. The proportion of amino acids mapped ar shown, light blue or light green is the P323 or D614 at the 323 positions in NSP12 and 614 in th spike protein, respectively. The proportion of the L or G in NSP12 and the spike protein respectively, is indicated in dark blue and dark green, respectively. The frequency of other amino acids at those positions is indicated in pink. We note that data were obtained through an ARTIC-Illumina based approach and as such PCR duplicates could not be removed. This may impact on the reported ratios.

Nasal washes were taken daily from each animal during infection ^13^. RNA was purified and sequenced using two independent approaches, shotgun sequencing on an Illumina platform and via ARTIC-Illumina with the latter for specifically sequencing SARS-CoV-2 RNA. Dominant viral genome sequence and minor genomic variants were determined for SARS-CoV-2 for each sample in which genome coverage could be obtained. To obtain a global overview and identify whether there were any hot spots for minor genomic variants, these were plotted as an average over the course of the infections in the non-human primates (NHPs) (Figure 4). The data indicated that minor genomic variants occurred throughout the genome, but the greatest variation occurred at position 14,408 in the orf1ab region, which resulted in a C to U change. This resulted in a non-synonymous change in NSP12 with the substitution of P323L (amino acid position 4715 with respect to the ORF1AB polyprotein).

**Figure 4.**
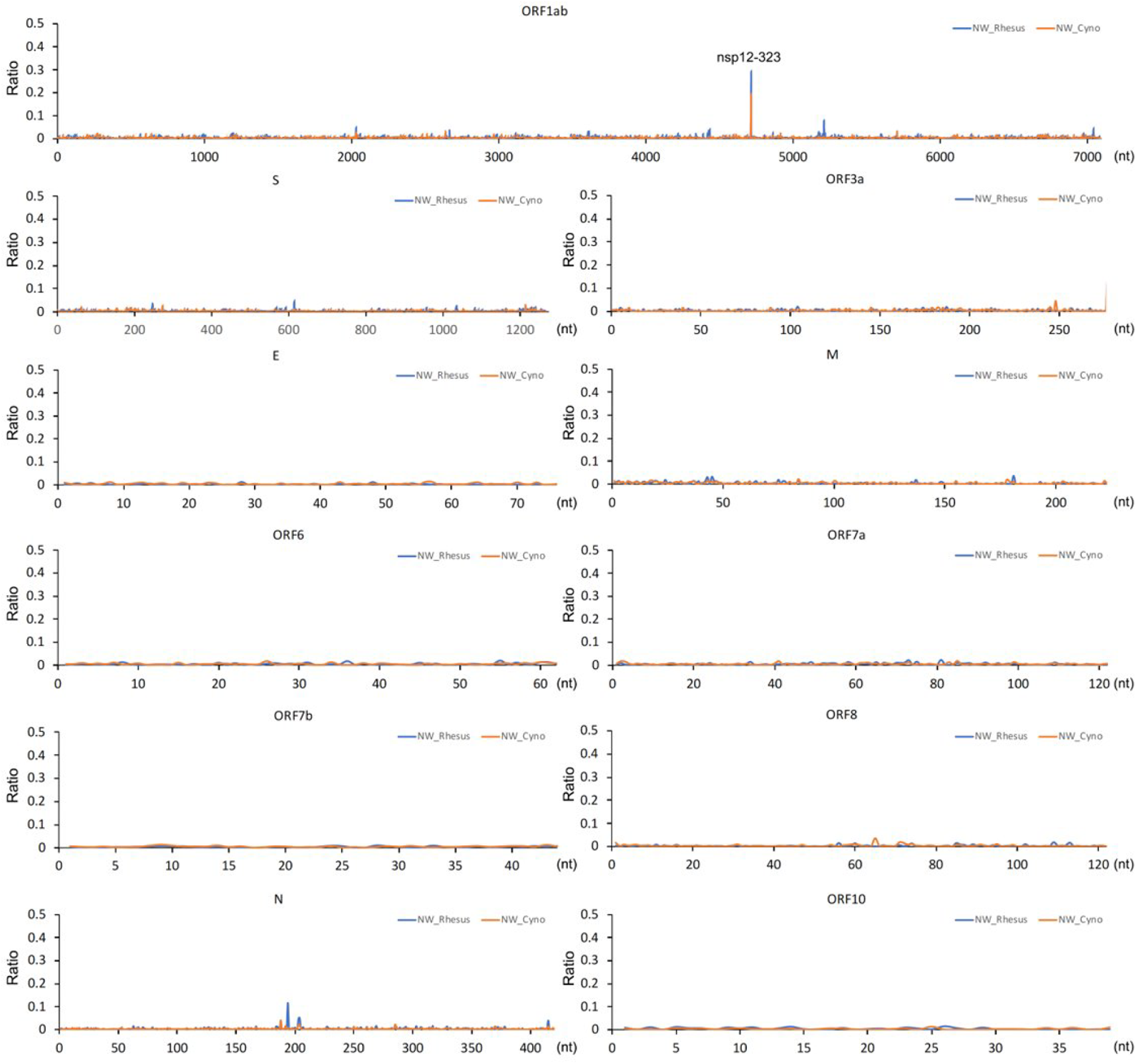
Analysis of minor variant genomes in cynomolgus and rhesus macaques infected with the SARS-CoV-2 Victoria/01/202040 isolate using data from shotgun Illumina RNA sequencing of nasal washes (NW). Data presented as a global average over the course of the infection from sequencing SARS-CoV-2 from longitudinal samples. Each SARS-CoV-2 open reading frame is indicated above the appropriate panel. The major difference was at position 323 in NSP12.

To determine how rapidly these mutations were selected in the individual animals, sequences from longitudinal samples were analyzed (Figure 5, showing ARTIC-Illumina data) (Supplementary Figure 1, showing both ARTIC-Illumina and ARTIC-Nanopore approaches and coverage). The sequencing data, using the two different approaches, showed that the P323L mutation was already present as a minor genomic variant (at higher levels than the inoculum) by Day 1 in some animals, as well as the presence of other minor genomic variants at this position. However, as infection progressed the frequency of the 323L minor genomic variant increased and became part of the dominant viral genome sequence by the end point of infection. This was the general pattern for all individual animals whether cynomolgus or rhesus macaque.

**Figure 5.**
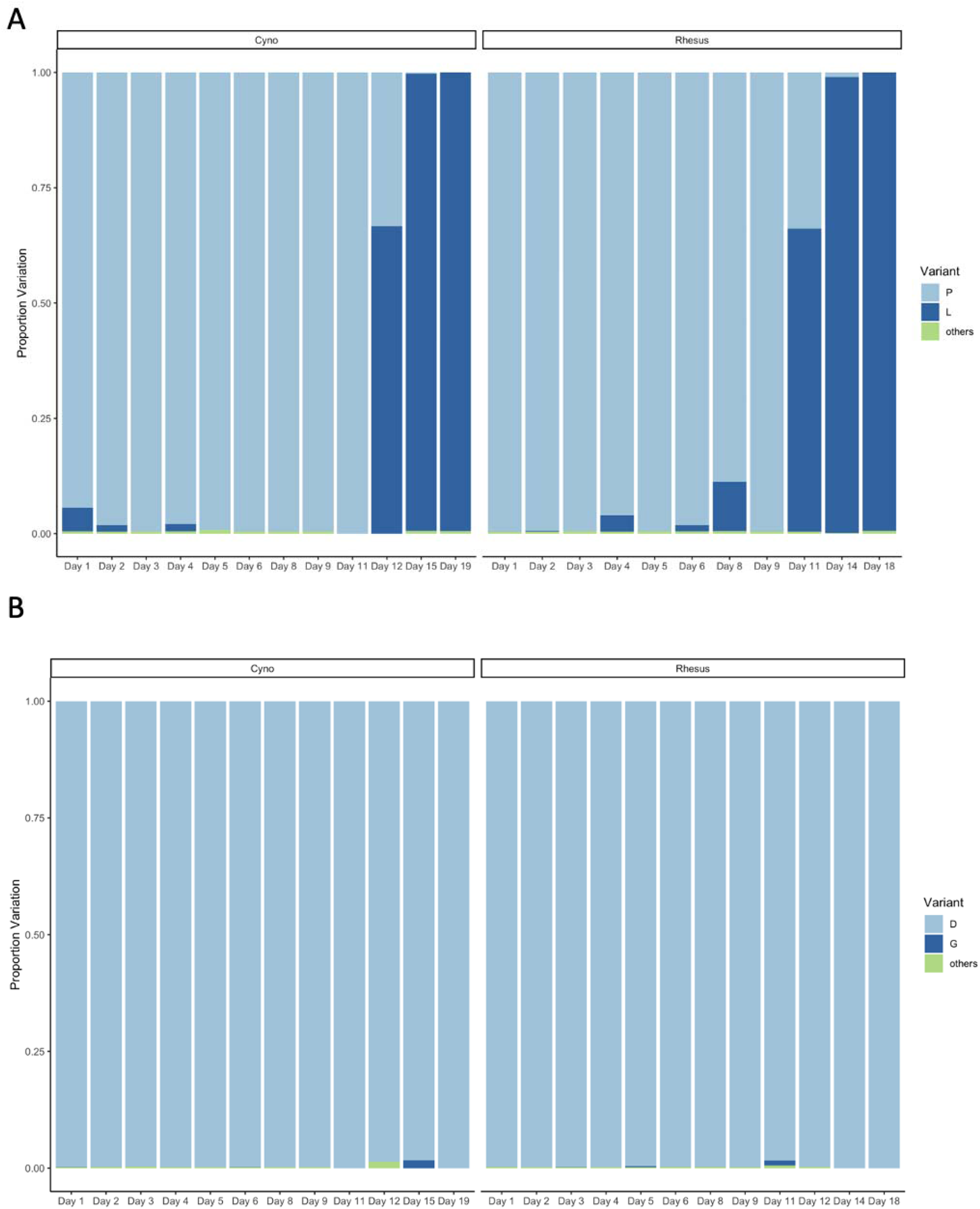
Analysis of NSP12 position 323 (A) and the spike protein position 614 (B) in SARS-CoV-2 from nasopharyngeal swabs taken longitudinally from infected cynomolgus and rhesus macaques. Data in this figure is from the ARTIC-Illumina approach to specifically amplify SARS-CoV-2 RNA. The day post infection is shown for the animals. In some cases, where there was more than one animal for each day, or usable sequence was obtained, the average value was calculated. For each position of interest either the P (for position 323 in NSP12) or D (for position 614 in the spike protein) is shown in light blue, and the substitution of L or G, shown in dark blue, respectively. Green indicates other substitutions at that position. The left-hand y-axis indicates the % variation at the indicated position). The % variation was only shown for these sites with coverage > 5.

### The P323L substitution in NSP12 confers a growth advantage in the context of a recombinant virus with 614G in the spike protein

Previous data indicated that Victoria/01/202040 grew with a small plaque phenotype and lower titer compared to more contemporary variants including Variants of Concern (VOCs), that grew to higher titres with larger or mixed plaque morphologies ^17^. The later virus isolates contained the P323L and D614G substitutions in NSP12 and the spike protein, respectively, as the dominant viral genome sequence, as well as other changes. To investigate whether the 323L substitution conferred an advantage over and above the 614G change in the spike protein, two recombinant viruses were created that were based on the 614G background, one with P323 (Wuhan/614G/P323) and the other with 323L (Wuhan/614G/323L) in NSP12. Growth of these two recombinant viruses were compared in cell culture by examining plaque morphology. The data indicated that Wuhan/614G/323L had a large plaque phenotype whereas Wuhan/614G/P323 had a small plaque phenotype (Figure 6), suggesting that the 323L substitution conferred a growth advantage.

**Figure 6.**
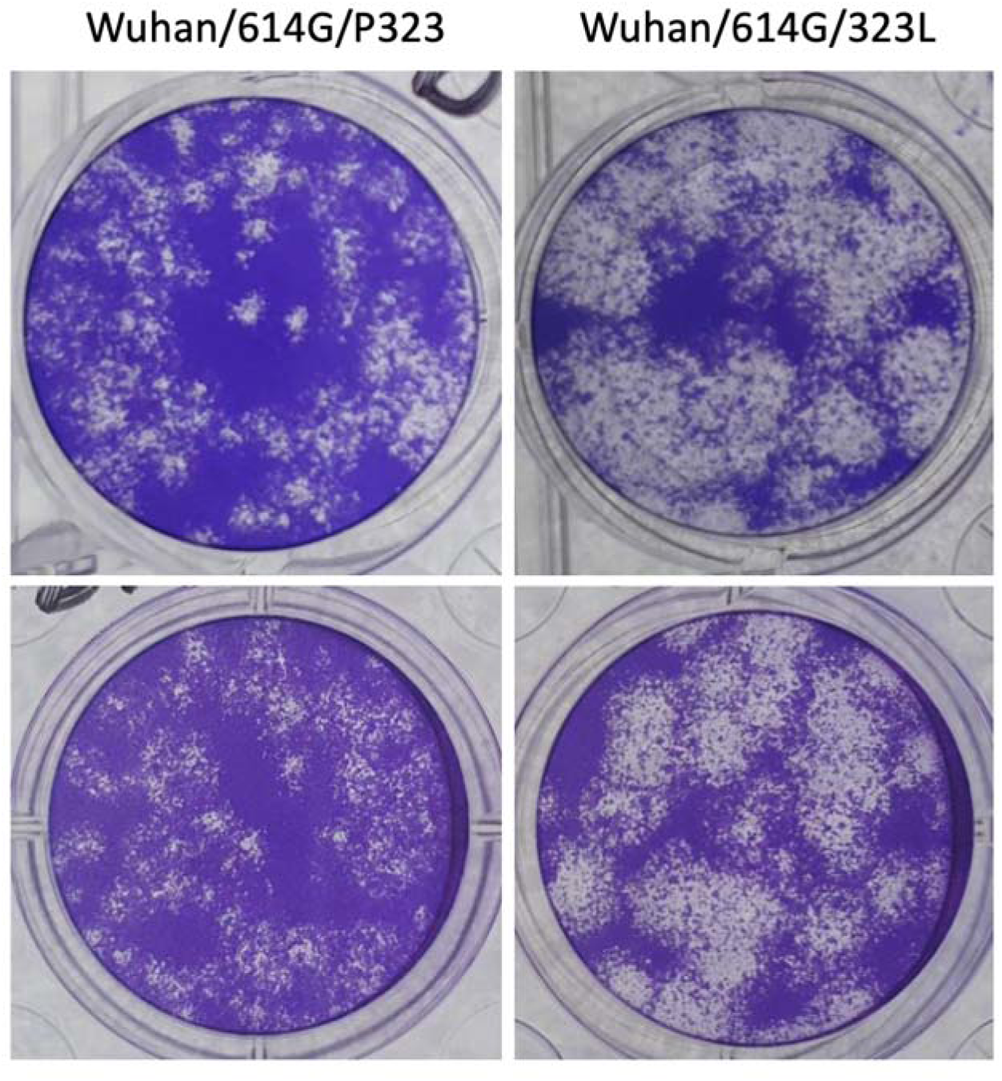
Representative images of plaques formed by two recombinant viruses that have the Wuhan-Hu-1 background (NC_045512) and an engineered D614G substitution in the spik protein and differed at position 323 in NSP12 with either a P or L, these were terme Wuhan/614G/P323 and Wuhan/614G/323L, respectively.

### Maintenance of variation at position 323 in NSP12 in the population

Based on the experimental data presented in this study, we propose a model where th emergence and distribution of minor variant genomes and dominant viral genome sequence for SARS-CoV-2 is dependent on selection pressure and time post-infection at which a virus population is transmitted onwards to another individual (Figure 7).

**Figure 7.**
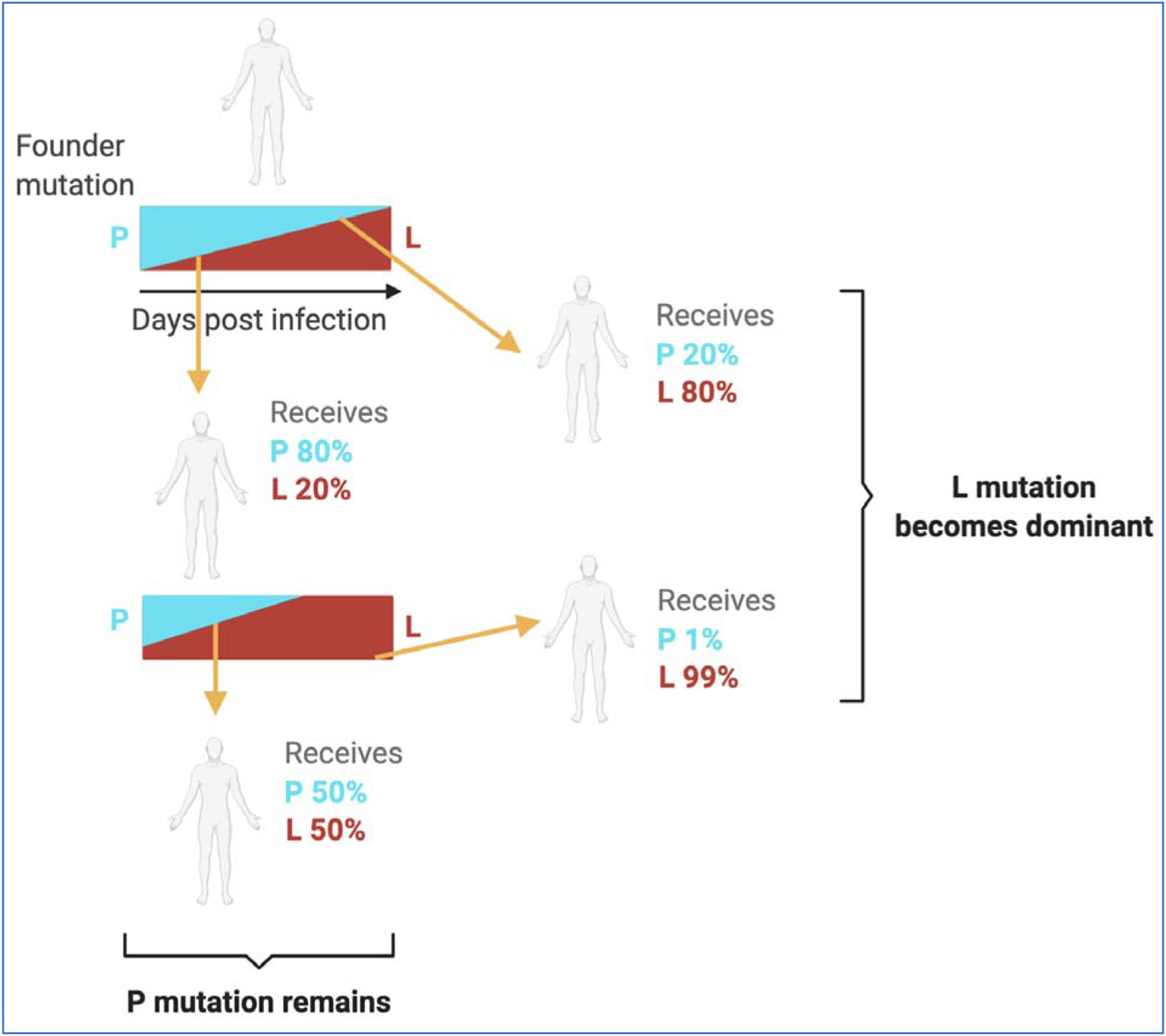
Model for the transmission of variant genomes which encodes amino acids under strong selection pressure showing the potential options for growth and transmission of viral populations with either consensus viral genomes with P323 (cyan) and 323L (red) present in minor variant genomes or in equilibrium or where 323L is in dominant viral genome sequence and P323 is present in the minor variant genomes. Given the potential strong selection pressure on this position the time post-infection transmission occurs is crucial in determining which variant becomes dominant viral genome sequence.

One of the predictions of this model is that whilst 323L in NSP12 might now be part of the dominant viral genome sequence, other variants at this position will be present and persist (e.g. P323) at this position. To test this contemporary sequence data (post the P323L and D614G substitutions) that had been deposited between July and September 2021 on the Short Read Archive was examined for variation at position 323 in NSP12 (Figure 8). The data indicated that 323L is the dominant variant, but P323 and other substitutions such as 323F are present as minor genomic variants.

**Figure 8.**
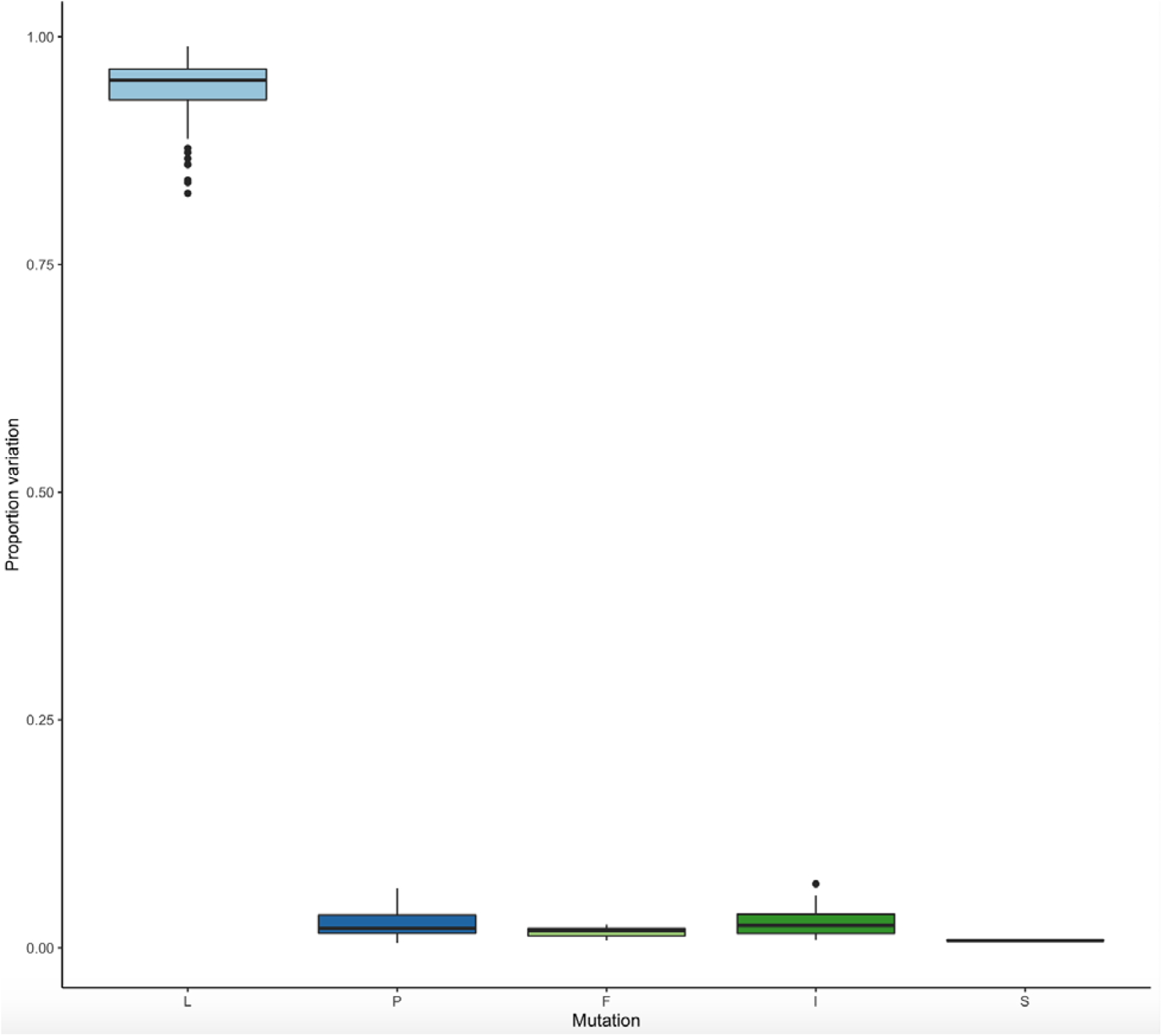
Amino acid mutations at site 323 in NSP12 in samples sequenced using the ARTIC-Nanopore approach (n=101) from July-September 2021 obtained from the Short Read Archive. The bioinformatics tool DiversiTools was used to generate proportions of the counts of amino acids at site 323 and showed that L is dominant in viral sequences from mid-late 2021, with P remaining a small proportion of the population alongside amino acids F, S and I.

## Discussion

Several variants have come to dominate the global landscape of SARS-CoV-2 infections, including ones with the initial D614G and P323L polymorphisms in the spike protein and NSP12 respectively (B.1), followed by Alpha (B.1.1.7), Delta (B.1.617.2) and Omicron (B.1.1529). These have occurred in waves and are likely linked to increases in transmissibility ^4^, coupled with spike variation-mediated immune escape ^18, 19^, founder effects ^20–22^, behaviour patterns of hosts and population density ^23, 24^ and non-pharmaceutical interventions ^25^. Whilst VoCs have differed in terms of transmissibility, in general there has been no marked change in inherent morbidity and mortality, although an early variant with a deletion in ORF8 was associated with a less severe inflammatory response and better patient outcome ^3^.

Among the first major changes in the dominant viral genome sequence of SARS-CoV-2 were the P323L and the D614G substitutions in NSP12 and the spike protein, respectively. Focus has been placed on spike D614G and its association with increased infectivity ^26^. We wanted to investigate the selection pressure at these two sites by analysing the virus population in humans over the period when the two substitutions became part of the dominant viral genome sequence, as well as studying this in two non-human primate animal models. The first analysis suggested rapid selection of P323L in NSP12 and D614G in the spike protein within humans. This was reflected in the substitutions 323L and 614G polymorphisms in the minor genomic variant population becoming the dominant viral genome sequence and replacing P323 and D614 within a few days of within host selection (Figure 2). At the population level, data suggested this selection was established over a two-month period in the UK (February and March 2020). We note that although samples used in this study were collected early in the pandemic in the UK, during the containment phase and in the early surge phase of Wave 1, there was no evidence that the change from P323L in NSP12 and D614G in the spike protein resulted in an increase in disease severity.

The selection pressure at these two positions (within an isolate close to the original Wuhan outbreak) was evaluated in two non-human primate models for COVID-19 that recapitulate the mild disease observed in most humans ^13^. Here, the SARS-CoV-2 variant used for infection had P323 in NSP12 and D614 in the spike protein in the dominant consensus sequence. At the minor variant genome level, 323L in NSP12 was present with a frequency of 0.03% and 614G in the spike protein at 0.02%. The sequence analysis indicated that for those animals where later time points returned usable viral genomic information, the dominant viral genome sequence now contained 323L in NSP12, but not necessarily 614G in the spike protein (Figure 5).

Recombinant viruses that differed at codon 323 in NSP12 in the context of a background with D614G in the spike protein and showed that the P323 virus grew with a smaller plaque morphology than a version with 323L. There are several different determinants of plaque size including those related to in vitro growth rate, evasion of antiviral responses and cell to cell Fusion ^27, 28^. NSP12 has been shown to attenuate type I interferon production ^29^, and this may be variant dependent. The mechanism behind the selection pressure acting on the P323L substitution in both humans and non-human primate animal models is unknown. However, NSP12 is the RNA dependent RNA polymerase, and such polymerase complexes can be composed of both viral and host cell proteins ^30, 31^. We speculate that the P323L substitution may alter the composition of the replication complex by altering interactions with the host cell proteome and thereby facilitating virus replication. Therefore, it is tempting to speculate that growth of viruses in cell lines from the original host species might drive the selection back. This might provide a mechanism to narrow down candidates for the original zoonotic event(s).

In our model (Figure 7), an individual with the substitution present in a minor variant genome with a selective advantage will see an increase in the proportion of this genome as infection progresses. Under this pressure the minor variant genome will become the dominant viral genome sequence. If transmission occurs early in infection, then the variant will be maintained at a minor genomic variant level. If selective pressure is strong then the viral population that is being transmitted will have the substitution as part of the dominant viral genome sequence – and this will persist during further infections. Another consequence is that the sudden emergence of a substitution as part of the dominant genome sequence may be due to founder effect. For example, 323F in NSP12 that was identified in a cluster of cases in Norther Nevada and in Nigeria (B.1.525). However, this substitution has not become part of the global dominant viral genome sequence, despite that 323F was identified in samples from early 2020.

The data in this study indicates that in some cases it may be possible to predict the emergence of a new dominant viral genome sequence and hence new variant. This would be based on tracking the distribution and frequency of minor variant genomes at a population level, rather than just focusing on providing information on the dominant viral genome sequence e.g., consensus level reporting. Whilst computationally more intensive and perhaps requiring higher quality samples and sequencing data, the ability to earlier predict a newly emerging variant of SARS-CoV-2 in the global landscape may aid in the evaluation of medical countermeasures and non-pharmaceutical interventions.

## Materials and methods

### Illumina for NHP NW samples

Total RNA in each sample was extracted with QIAmp viral RNA extraction kit and eluted in pure water. Following the manufacturer’s protocols, total RNA was used as input material in to the QIAseq FastSelect –rRNA HMR (Qiagen) protocol to remove cytoplasmic and mitochondrial rRNA with a fragmentation time of 7 or 15 minutes. Subsequently, the NEBNext® Ultra™ II Directional RNA Library Prep Kit for Illumina® (New England Biolabs) was used to generate the RNA libraries, followed by 11 cycles of amplification and purification using AMPure XP beads. Each library was quantified using Qubit and the size distribution assessed using the Agilent 2100 Bioanalyser, and the final libraries were pooled in equimolar ratios. The raw FASTQ files (2 x 150 bp) generated by an Illumina® NovaSeq 6000 (Illumina®, San Diego, USA) were trimmed to remove Illumina adapter sequences using Cutadapt v1.2.1 ^32^. The option “−O 3” was set, so the that 3’ end of any reads which matched the adapter sequence with greater than 3 bp was trimmed off. The reads were further trimmed to remove low quality bases, using Sickle v1.200 ^33^ with a minimum window quality score of 20. After trimming, reads shorter than 10 bp were removed.

The minor variations of amino acid in the genes of virus were called as our previous description ^34^. Hisat2 v2.1.0 ^35^ was used to map the trimmed reads on the cynomolgus (*M. fascicularis*) and rhesus (*M. mulatta*) reference genome assemblies (release-94) downloaded from the Ensembl FTP site. The unmapped reads were extracted by bam2fastq (v1.1.0) and then mapped on the inoculum SARS-CoV-2 genome (GenBank sequence accession: NC_045512.2) using Bowtie2 v2.3.5.1 ^35^ by setting the options to parameters “--local -X 2000 --no-mixed”, followed by SAM file to BAM file conversion, sorting, and removal of the reads with a mapping quality score below 11 using SAMtools v1.9 ^36^. After that, the PCR and optical duplicate reads in the BAM files were discarded using the MarkDuplicates in the Picard toolkit v2.18.25 (http://broadinstitute.github.io/picard/) with the option of “REMOVE_DUPLICATES=true”. This BAM file was then processed by the diversiutils script in DiversiTools (http://josephhughes.github.io/btctools/) with the “-orfs” function to generate the number of amino acid changes caused by the nucleotide deviation at each site in the protein. In order to distinguish low frequency variants from Illumina sequence errors, the diversiutils script used the calling algorithms based on the Illumina quality scores to calculate a P-value for each variant at each nucleotide site ^37^. The amino acid change was then filtered based on the P-value (<0.05) to remove the low frequency variants from Illumina sequence errors.

### ARTIC Illumina for longitudinal swab samples and NHP NW samples

Samples from clinical specimens were processed at CL3 at the University of Liverpool as part of the study described in this chapter. Nasopharyngeal swabs were collected in viral transport media. Swabs were left to defrost in a Tripass I cabinet in CL3. The swab was removed from the tube and dipped in virkon before disposal to reduce dripping and aerosol generation. 250ml of viral transport media was removed from the swab sample and added to 750ml of Trizol LS (Invitrogen (10296028)) and mixed well. Remaining extraction was continued under CL2 conditions. All RNA samples were then treated with Turbo DNase (Invitrogen). SuperScript IV (Invitrogen) was used to generate single-strand cDNA using random primer mix (NEB, Hitchin, UK). ARTIC V3 PCR amplicons from the single-strand cDNA were generated following the Nanopore Protocol of PCR tiling of SARS-CoV-2 virus (Version: PTC_9096_v109_revL_06Feb2020). The amplicons products were then used in Illumina NEBNext Ultra II DNA Library preparation. Following 4 cycles of amplification the library was purified using Ampure XP beads and quantified using Qubit and the size distribution assessed using the Fragment Analyzer. Finally, the ARTIC library was sequenced on the Illumina® NovaSeq 6000 platform (Illumina®, San Diego, USA) following the standard workflow. The generated raw FASTQ files (2 x 250 bp) were trimmed to remove Illumina adapter sequences using Cutadapt v1.2.1 26. The option “−O 3” was set, so the that 3’ end of any reads which matched the adapter sequence with greater than 3 bp was trimmed off. The reads were further trimmed to remove low quality bases, using Sickle v1.200 27 with a minimum window quality score of 20. After trimming, reads shorter than 10 bp were removed. The NHP NW total RNA have been extracted and sequenced in our previous paper ^38^.

The variations of amino acid in the genes of the virus were called as our previous description ^34^. Hisat2 v2.1.0 ^35^ was used to map the trimmed reads onto the human reference genome assembly GRCh38 (release-91) downloaded from the Ensembl FTP site. The unmapped reads were extracted by bam2fastq (v1.1.0) and then mapped on a known SARS-CoV-2 genome (GenBank sequence accession: NC_045512.2) using Bowtie2 v2.3.5.1 ^35^ by setting the options to parameters “--local -X 500 --no-mixed”, followed by SAM file to BAM file conversion, sorting, and removal of the reads with a mapping quality score below 11, not in pair, and not primary and supplementary alignment using SAMtools v1.9 ^36^. Bamclipper (v 1.0.0) ^39^ was used to trim the ARTIC primer sequences on the mapped reads within the BAM files. The reads without ARTIC primer sequences were also excluded in the further analysis. This trimmed BAM file was then processed by the diversiutils script in DiversiTools (http://josephhughes.github.io/DiversiTools/) with the “-orfs” function to generate the number of amino acid changes caused by the nucleotide deviation at each site in the protein in comparison to the reference SARS-CoV-2 genome (NC_045512.2). In order to distinguish low frequency variants from Illumina sequence errors, the diversiutils script used the calling algorithms based on the Illumina quality scores to calculate a P-value for each variant at each nucleotide site ^37^.

### Rapid Sequencing Long Amplicons (RSLA) nanopore for longitudinal swab samples

Total RNA of longitudinal swab samples were extracted as described above. Sequencing libraries for amplicons generated by RSLA ^14^ were prepared following the ‘PCR tiling of SARS-CoV-2 virus with Native Barcoding’ protocol provided by Oxford Nanopore Technologies using LSK109 and EXP-NBD104/114. The artic-ncov2019 pipeline v1.2.1 (https://artic.network/ncov-2019/ncov2019-bioinformatics-sop.html) was used to filter the passed FASTQ files produced by Nanopore sequencing with lengths between 800 and 1600. This pipeline was then used to map the filtered reads on the reference SARS-CoV-2 genome (NC_045512.2) by minimap2 and assigned each read alignment to a derived amplicon and excluded primer sequences based on the RSLA primer schemes in the BAM files. These BAM files were further analysed using DiversiTools (http://josephhughes.github.io/btctools/) with the “-orfs” function to generate the ratio of amino acid change in the reads and coverage at each site of the protein in comparison to the reference SARS-CoV-2 genome (NC_045512.2). The amino acids with highest ratio and coverage > 10 were used to assemble the consensus protein sequences.

### Sanger sequencing

cDNA template was amplified using Q5 High-Fidelity DNA Polymerase following the PCR conditions: denaturation at 98°C for 30 sec followed by 39 cycles of 10 sec denaturation at 98°C, 30 sec annealing at 66°C, and then 50 sec of extension at 72°C. A final extension step was done for 2 min at 72°C. The primer sets used for amplification were (SARS-CoV-2_15_LEFT=ATACGCCAACTTAGGTGAACG, SARS-CoV-2_15_RIGHT= AACATGTTG-TGCCAACCACC) to detect the P323L mutation or (SARS-CoV-2_24_LEFT= TTGAACTTCTACATGCACCAGC, SARS-CoV-2_RIGHT=CCAGAAGTGATTGTACCCGC) to detect the D614G mutation. PCR products were purified using AMPure XP beads (Beckman Coulter) and quantified using the Qubit High Sensitivity 1X dsDNA kit (Invitrogen). To visualise band quality, PCR products were run on a 1.5% agarose gel. 10 ng of each amplified product was sent for sanger sequencing (Source Bioscience, UK).

### Cells

African green monkey kidney C1008 (Vero E6) cells (Public Health England, PHE) were cultured in Dulbecco’s minimal essential medium (DMEM) (Sigma) with 10% foetal bovine serum (FBS) (Sigma) and 0.05mg/ml gentamicin at 37°C/5% CO2. Vero/hSLAM cells (PHE) were grown in DMEM with 10% FBS and 0.05mg/ml gentamicin (Merck) with the addition of 0.4mg/ml Geneticin (G418; Thermofisher) at 37°C/5% CO2. Human ACE2-A549 (hACE2-A549), a lung epithelial cell line which overexpresses the ACE-2 receptor ^40^, were cultured in DMEM with 10% FBS and 0.05mg/ml gentamicin with the addition of 10µg/ml Blasticidin (Invitrogen). Only passage 3-10 cultures were used for experiments.

### Generation and culture of recombinant viruses

Recombinant SARS-CoV-2 viruses were generated by reverse genetics using the “transformation-associated recombination” in yeast approach ^41^. 11 cDNA fragments with 70 bp end-terminal overlaps which spanned the entire SARS-CoV-2 isolate Wuhan-Hu-1 genome (GenBank accession: NC_045512) were produced by GeneArt™ synthesis (Invitrogen™, ThermoFisher) as inserts in sequence verified, stable plasmid clones. The 5′ terminal cDNA fragment was modified to contain a T7 RNA polymerase promoter and an extra “G” nucleotide immediately upstream of the SARS-CoV-2 5′ sequence, whilst the 3′ terminal cDNA fragment was modified such that the 3’ end of the SARS-CoV-2 genome was followed by a stretch of 33 “A”s followed by the unique restriction enzyme site Asc I. The inserts were amplified by PCR using a Platinum SuperFi II mastermix (ThermoFisher) and assembled into full length SARS-CoV-2 cDNA clones in the YAC vector pYESL1 using a GeneArt™ High-Order Genetic Assembly System (A13285, Invitrogen™, ThermoFisher) according to the manufacturer’s instructions. RNA transcripts produced from the YAC clones by transcription with T7 polymerase were used to recover infectious virus. Two viruses were produced on the Wuhan-Hu-1 background and had a D614G substitution in the spike protein and differed at amino acid position 323 in NSP12 with either a P or L, these were termed Wuhan/614G/P323 and Wuhan/614G/323L, respectively. Whole genome sequencing confirmed the presence of these changes. Stocks of the viruses were cultured in Vero E6 cells in DMEM containing 2% FBS, 0.05mg/ml gentamicin and harvested 72 hours post inoculation. Virus stocks were aliquoted and stored at −80°C. All stocks were titred by plaque assay on Vero E6 cells and pictures of the resulting plaques recorded.

### Serial passage of SARS-CoV-2 Victoria/01/2020

SARS-CoV-2 Victoria/01/2020 was passaged three times in Vero/hSLAM cells prior to receiving it. hACE2-A549 cells were then infected at an MOI of 0.01 and incubated for 72 hours (Passage 4). Following this, 100μl was passaged to fresh cells and incubated at 37C for 1 hour. After the incubation, media was topped up with DMEM containing 2% FBS, 0.05mg/ml gentamicin and incubated for 72 hours (Passage 5). This process was repeated until Passage 13 (a total of ten passages through hACE2-A549 cells).

### Analysis of global sequences from July-September 2021

Sequences were obtained from the Short Read Archive (SRA) under accession numbers: ERR6343731, ERR6343734, ERR6343745, ERR6343747, ERR6343749, ERR6344225, ERR6346453, ERR6346456, ERR6346459, ERR6758978, ERR6758981, ERR6759296, ERR6761288, ERR6761458, ERR6761562, ERR6761570, ERR6761711, ERR6761986, ERR6762387, ERR6762545, ERR6762546, ERR6825821, ERR6878898, ERR6879599, ERR6879604, ERR6887797, ERR6887811, ERR6887812, ERR6887820, ERR6888048, ERR6888063, ERR6888078, ERR6888265, ERR6888283, SRR16376487, SRR16376490, SRR16376491, SRR16376494, SRR16376495, SRR16376496, SRR16376497, SRR16376501, SRR16376502, SRR16376505, SRR16376510, SRR16376515, SRR16376516, SRR16376522, SRR16376523, SRR16376524, SRR16376526, SRR16376529, SRR16376530, SRR16376531, SRR16376536, SRR16376540, SRR16376543, SRR16376544, SRR16376547, SRR16376551, SRR16376552, SRR16376554, SRR16376557, SRR16376559, SRR16376573, SRR16376580, SRR16376589, SRR16376599, SRR16376608, SRR16376613, SRR16376614, SRR16376648, SRR16376678, SRR16376782, SRR16376802, SRR16376804, SRR16376807, SRR16376810, SRR16376884, SRR16376904, SRR16376907, SRR16376912, SRR16376913, SRR16376914, SRR16376916, SRR16376921, SRR16376922, SRR16376925, SRR16376927, SRR16376928, SRR16376929, SRR16376932, SRR16376935, SRR16376939, SRR16376940, SRR16376941, SRR16376943, SRR16376944, SRR16376946, SRR16376949, SRR16376951. All sequences were ARTIC-Nanopore sequenced using the V3 primer scheme and downloaded as SRA files. The SRA files were converted to FASTQ files using the SRA Toolkit v2.11.3 (https://github.com/ncbi/sra-tools) command fastq-dump. The FASTQ files were processed through the artic-ncov2019 v1.2.1 pipeline (https://artic.network/ncov-2019/ncov2019-bioinformatics-sop.html) and the DiversiTools tool (https://github.com/josephhughes/DiversiTools) as described above.

## Acknowledgments

We would like to thank all members of the Hiscox Laboratory and the Centre for Genome Research for supporting SARS-CoV-2/COVID-19 sequencing research and members of ISARIC4C consortia (see supplementary information and https://isaric4c.net/about/authors/). The authors would like to thank J. Druce and M.G. Catton from the Victorian Infectious Diseases Reference Laboratory, Royal Melbourne Hospital, at the Peter Doherty Institute for Infection and Immunity, Victoria, 3000, Australia, for providing the SARS-CoV-2 isolate used in this study. We would like to thank Oliver Schwartz for the gift of ACE2-A549 cells. This work was funded by U.S. Food and Drug Administration Medical Countermeasures Initiative contract (75F40120C00085) to JAH with Co-Is, MWC, ADD, AD, DAM, MGS and LT. The article reflects the views of the authors and does not represent the views or policies of the FDA. The non-human primate work was funded by the Coalition of Epidemic Preparedness Innovations (CEPI) and the Medical Research Council Project CV220-060, “Development of an NHP model of infection and ADE with COVID-19 (SARS-CoV-2) both awarded to MWC. This work was also supported by the MRC (MR/W005611/1) G2P-UK: A national virology consortium to address phenotypic consequences of SARS-CoV-2 genomic variation (co-Is ADD and JAH). JAH is also funded by the Centre of Excellence in Infectious Diseases Research (CEIDR) and the Alder Hey Charity. The ISARIC4C sample collection and sequencing in this study was supported by grants from the Medical Research Council (grant MC_PC_19059), the National Institute for Health Research (NIHR; award CO-CIN-01) and the Medical Research Council (MRC; grant MC_PC_19059). JAH, MGS, MWC and LT are supported by the NIHR Health Protection Research Unit (HPRU) in Emerging and Zoonotic Infections at University of Liverpool in partnership with the UK Health Security Agency (UK-HSA), in collaboration with Liverpool School of Tropical Medicine and the University of Oxford (award 200907). LT is supported by a Wellcome Trust fellowship [205228/Z/16/Z]. PD and JKB acknowledge Institute Strategic Programme grant (no. BB/P013740/1) from the BBSRC.

## Author contributions

Conceptualization: DAM, AD, MWC and JAH. Data curation: HG, XD, RP-R, DAM, AD and JAH. Formal analysis: HG, XD, NR, RP-R, PD, ADD, TP, AD and JAH. Funding acquisition: MGS, PJMO, JKB, DAM, LT, AD, ADD, MWC and JAH. Investigation: HG, XD, NR, RP-R, ADD, GTS, BJ, MKW, ME, JB, TJ, FJS, SRE, JT, CH and JAH. Methodology: HG, XD, GTS, RP-R, ADD, MKW, FJS and JT. Project administration: JAH. Resources: MWC, FJS, JT, YH, MGS, PJMO, ADD, JKB, LT. Software: HG, XD, RP-R, DAM and AD. Supervision: MWC, AD, ADD, SRE and JAH. Validation: HG, XD, RP-R, NR, CH, GTS, DAM, AD and JAH. Visualisation: HG, XD and RP-R. Writing – original draft: HG, XD, RP-R and JAH. Writing – reviewing and editing. HG, XD, RP-R, DAM, AD, LT, ADD, MWC and JAH.

## Availability of data and materials

All viral sequence data used in this analysis were deposited with the National Center for Biotechnology Information under the project accession number PRJNA789459 and can be accessed via https://www.ncbi.nlm.nih.gov/bioproject/PRJNA789459.

## Competing interests

The authors declare that they have no competing interests.

## Ethics approval and consent to participate

Patients were recruited under the International Severe Acute Respiratory and emerging Infection Consortium (ISARIC) Clinical Characterisation Protocol CCP (https://isaric.net/ccp) by giving informed consent. ISARIC CCP was reviewed and approved by the national research ethics service, Oxford (13/SC/0149). All experimental work on non-human primates was conducted under the authority of a UK Home Office approved project license (PDC57C033) that had been subject to local ethical review at PHE Porton Down by the Animal Welfare and Ethical Review Body (AWERB) and approved as required by the Home Office Animals (Scientific Procedures) Act 1986. None of the animals had been used previously for experimental procedures.

## Supplementary information

**Supplementary Figure 1.**
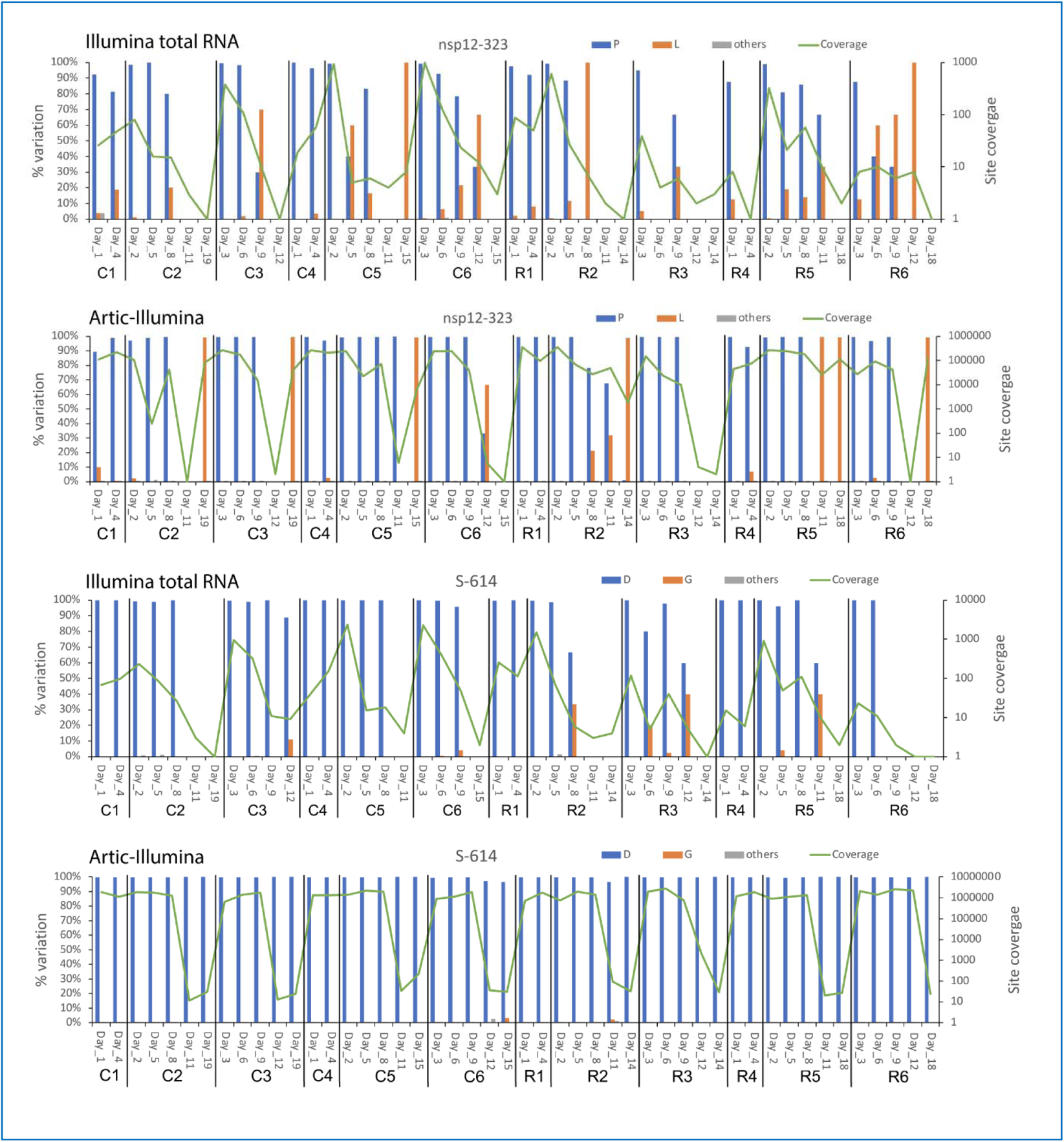
Analysis of NSP12 position 323 and the spike protein position 614 i SARS-CoV-2 from nasopharyngeal swabs taken longitudinally from infected cynomolgus (C) and rhesus (R) macaques. Sequencing was performed using both an Illumina shot gun sequencing approach (Illumina total RNA) or using an ARTIC-Illumina approach to specifically amplify SARS-CoV-2 RNA. The day post infection is shown for each individual animal (number after the C or R) (x-axis). For each position of interest either the P (for position 323 in NSP12) or D (for position 614 in the spike protein) is shown in blue, and the substitution of L or G, shown in orange, respectively. Grey indicates other substitutions at that position. The left-hand y-axis indicates the % variation at the indicated position and the right-hand x-axis shows amino acid site coverage for each position (green line). The % variation was only shown for these sites with coverage > 5.

## Supplementary information

ISARIC4C Investigators

Consortium Lead Investigator: J Kenneth Baillie.

Chief Investigator: Malcolm G Semple.

Co-Lead Investigator: Peter JM Openshaw.

ISARIC Clinical Coordinator: Gail Carson.

Co-Investigator: Beatrice Alex, Petros Andrikopoulos, Benjamin Bach, Wendy S Barclay, Debby Bogaert, Meera Chand, Kanta Chechi, Graham S Cooke, Ana da Silva Filipe, Thushan de Silva, Annemarie B Docherty, Gonçalo dos Santos Correia, Marc-Emmanuel Dumas, Jake Dunning, Tom Fletcher, Christoper A Green, William Greenhalf, Julian L Griffin, Rishi K Gupta, Ewen M Harrison, Julian A Hiscox, Antonia Ying Wai Ho, Peter W Horby, Samreen Ijaz, Saye Khoo, Paul Klenerman, Andrew Law, Matthew R Lewis, Sonia Liggi, Wei Shen Lim, Lynn Maslen, Alexander J Mentzer, Laura Merson, Alison M Meynert, Shona C Moore, Mahdad Noursadeghi, Michael Olanipekun, Anthonia Osagie, Massimo Palmarini, Carlo Palmieri, William A Paxton, Georgios Pollakis, Nicholas Price, Andrew Rambaut, David L Robertson, Clark D Russell, Vanessa Sancho-Shimizu, Caroline J Sands, Janet T Scott, Louise Sigfrid, Tom Solomon, Shiranee Sriskandan, David Stuart, Charlotte Summers, Olivia V Swann, Zoltan Takats, Panteleimon Takis, Richard S Tedder, AA Roger Thompson, Emma C Thomson, Ryan S Thwaites, Lance CW Turtle, Maria Zambon.

Project Manager: Hayley Hardwick, Chloe Donohue, Fiona Griffiths, Wilna Oosthuyzen.

Project Administrator: Cara Donegan, Rebecca G. Spencer.

Data Analyst: Lisa Norman, Riinu Pius, Thomas M Drake, Cameron J Fairfield, Stephen R Knight, Kenneth A Mclean, Derek Murphy, Catherine A Shaw.

Data and Information System Manager: Jo Dalton, Michelle Girvan, Egle Saviciute, Stephanie Roberts, Janet Harrison, Laura Marsh, Marie Connor, Sophie Halpin, Clare Jackson, Carrol Gamble, Daniel Plotkin, James Lee.

Data Integration and Presentation: Gary Leeming, Andrew Law, Murray Wham, Sara Clohisey, Ross Hendry, James Scott-Brown.

Material Management: Victoria Shaw, Sarah E McDonald.

Patient Engagement: Seán Keating.

Outbreak Laboratory Staff and Volunteers: Katie A. Ahmed, Jane A Armstrong, Milton Ashworth, Innocent G Asiimwe, Siddharth Bakshi, Samantha L Barlow, Laura Booth, Benjamin Brennan, Katie Bullock, Benjamin WA Catterall, Jordan J Clark, Emily A Clarke, Sarah Cole, Louise Cooper, Helen Cox, Christopher Davis, Oslem Dincarslan, Chris Dunn, Philip Dyer, Angela Elliott, Anthony Evans, Lorna Finch, Lewis WS Fisher, Terry Foster, Isabel Garcia-Dorival, Philip Gunning, Catherine Hartley, Rebecca L Jensen, Christopher B Jones, Trevor R Jones, Shadia Khandaker, Katharine King, Robyn T. Kiy, Chrysa Koukorava, Annette Lake, Suzannah Lant, Diane Latawiec, Lara Lavelle-Langham, Daniella Lefteri, Lauren Lett, Lucia A Livoti, Maria Mancini, Sarah McDonald, Laurence McEvoy, John McLauchlan, Soeren Metelmann, Nahida S Miah, Joanna Middleton, Joyce Mitchell, Shona C Moore, Ellen G Murphy, Rebekah Penrice-Randal, Jack Pilgrim, Tessa Prince, Will Reynolds, P. Matthew Ridley, Debby Sales, Victoria E Shaw, Rebecca K Shears, Benjamin Small, Krishanthi S Subramaniam, Agnieska Szemiel, Aislynn Taggart, Jolanta Tanianis-Hughes, Jordan Thomas, Erwan Trochu, Libby van Tonder, Eve Wilcock, J. Eunice Zhang, Lisa Flaherty, Nicole Maziere, Emily Cass, Alejandra Doce Carracedo, Nicola Carlucci, Anthony Holmes, Hannah Massey.

Edinburgh Laboratory Staff and Volunteers: Lee Murphy, Sarah McCafferty, Richard Clark, Angie Fawkes, Kirstie Morrice, Alan Maclean, Nicola Wrobel, Lorna Donnelly, Audrey Coutts, Katarzyna Hafezi, Louise MacGillivray, Tammy Gilchrist.

Local Principal Investigators: Kayode Adeniji, Daniel Agranoff, Ken Agwuh, Dhiraj Ail, Erin L. Aldera, Ana Alegria, Sam Allen, Brian Angus, Abdul Ashish, Dougal Atkinson, Shahedal Bari, Gavin Barlow, Stella Barnass, Nicholas Barrett, Christopher Bassford, Sneha Basude, David Baxter, Michael Beadsworth, Jolanta Bernatoniene, John Berridge, Colin Berry, Nicola Best, Pieter Bothma, David Chadwick, Robin Brittain-Long, Naomi Bulteel, Tom Burden, Andrew Burtenshaw, Vikki Caruth, David Chadwick, Duncan Chambler, Nigel Chee, Jenny Child, Srikanth Chukkambotla, Tom Clark, Paul Collini, Catherine Cosgrove, Jason Cupitt, Maria-Teresa Cutino-Moguel, Paul Dark, Chris Dawson, Samir Dervisevic, Phil Donnison, Sam Douthwaite, Andrew Drummond, Ingrid DuRand, Ahilanadan Dushianthan, Tristan Dyer, Cariad Evans, Chi Eziefula, Chrisopher Fegan, Adam Finn, Duncan Fullerton, Sanjeev Garg, Sanjeev Garg, Atul Garg, Effrossyni Gkrania-Klotsas, Jo Godden, Arthur Goldsmith, Clive Graham, Elaine Hardy, Stuart Hartshorn, Daniel Harvey, Peter Havalda, Daniel B Hawcutt, Maria Hobrok, Luke Hodgson, Anil Hormis, Michael Jacobs, Susan Jain, Paul Jennings, Agilan Kaliappan, Vidya Kasipandian, Stephen Kegg, Michael Kelsey, Jason Kendall, Caroline Kerrison, Ian Kerslake, Oliver Koch, Gouri Koduri, George Koshy, Shondipon Laha, Steven Laird, Susan Larkin, Tamas Leiner, Patrick Lillie, James Limb, Vanessa Linnett, Jeff Little, Mark Lyttle, Michael MacMahon, Emily MacNaughton, Ravish Mankregod, Huw Masson, Elijah Matovu, Katherine McCullough, Ruth McEwen, Manjula Meda, Gary Mills, Jane Minton, Mariyam Mirfenderesky, Kavya Mohandas, Quen Mok, James Moon, Elinoor Moore, Patrick Morgan, Craig Morris, Katherine Mortimore, Samuel Moses, Mbiye Mpenge, Rohinton Mulla, Michael Murphy, Megan Nagel, Thapas Nagarajan, Mark Nelson, Lillian Norris, Matthew K. O’Shea, Igor Otahal, Marlies Ostermann, Mark Pais, Carlo Palmieri, Selva Panchatsharam, Danai Papakonstantinou, Hassan Paraiso, Brij Patel, Natalie Pattison, Justin Pepperell, Mark Peters, Mandeep Phull, Stefania Pintus, Jagtur Singh Pooni, Tim Planche, Frank Post, David Price, Rachel Prout, Nikolas Rae, Henrik Reschreiter, Tim Reynolds, Neil Richardson, Mark Roberts, Devender Roberts, Alistair Rose, Guy Rousseau, Bobby Ruge, Brendan Ryan, Taranprit Saluja, Matthias L Schmid, Aarti Shah, Prad Shanmuga, Anil Sharma, Anna Shawcross, Jeremy Sizer, Manu Shankar-Hari, Richard Smith, Catherine Snelson, Nick Spittle, Nikki Staines, Tom Stambach, Richard Stewart, Pradeep Subudhi, Tamas Szakmany, Kate Tatham, Jo Thomas, Chris Thompson, Robert Thompson, Ascanio Tridente, Darell Tupper-Carey, Mary Twagira, Nick Vallotton, Rama Vancheeswaran, Lisa Vincent-Smith, Shico Visuvanathan, Alan Vuylsteke, Sam Waddy, Rachel Wake, Andrew Walden, Ingeborg Welters, Tony Whitehouse, Paul Whittaker, Ashley Whittington, Padmasayee Papineni, Meme Wijesinghe, Martin Williams, Lawrence Wilson, Sarah Cole, Stephen Winchester, Martin Wiselka, Adam Wolverson, Daniel G Wootton, Andrew Workman, Bryan Yates, Peter Young.

## Notes

### Competing Interest Statement

The authors have declared no competing interest.

## References

1. Worobey, M. et al. The emergence of SARS-CoV-2 in Europe and North America. Science 370, 564–570, doi:10.1126/science.abc8169 (2020).

2. Davidson, A. D. et al. Characterisation of the transcriptome and proteome of SARS-CoV-2 reveals a cell passage induced in-frame deletion of the furin-like cleavage site from the spike glycoprotein. Genome Med 12, 68, doi:10.1186/s13073-020-00763-0 (2020).

3. Young, B. E. et al. Effects of a major deletion in the SARS-CoV-2 genome on the severity of infection and the inflammatory response: an observational cohort study. Lancet 396, 603–611, doi:10.1016/S0140-6736(20)31757-8 (2020).

4. Hou, Y. J. et al. SARS-CoV-2 D614G variant exhibits efficient replication ex vivo and transmission in vivo. Science, doi:10.1126/science.abe8499 (2020).

5. Yang, H. C. et al. Analysis of genomic distributions of SARS-CoV-2 reveals a dominant strain type with strong allelic associations. Proc Natl Acad Sci U S A, doi:10.1073/pnas.2007840117 (2020).

6. Simmonds, P. Rampant C-->U Hypermutation in the Genomes of SARS-CoV-2 and Other Coronaviruses: Causes and Consequences for Their Short- and Long-Term Evolutionary Trajectories. mSphere 5, doi:10.1128/mSphere.00408-20 (2020).

7. Ratcliff, J. & Simmonds, P. Potential APOBEC-mediated RNA editing of the genomes of SARS-CoV-2 and other coronaviruses and its impact on their longer term evolution. Virology 556, 62–72, doi:10.1016/j.virol.2020.12.018 (2021).

8. Dong, X. et al. Identification and quantification of SARS-CoV-2 leader subgenomic mRNA gene junctions in nasopharyngeal samples shows phasic transcription in animal models of COVID-19 and dysregulation at later time points that can also be identified in humans. bioRxiv, 2021.2003.2003.433753, doi:10.1101/2021.03.03.433753 (2021).

9. Peacock, T. P., Penrice-Randal, R., Hiscox, J. A. & Barclay, W. S. SARS-CoV-2 one year on: evidence for ongoing viral adaptation. J Gen Virol 102, doi:10.1099/jgv.0.001584 (2021).

10. Lythgoe, K. A. et al. SARS-CoV-2 within-host diversity and transmission. Science 372, doi:10.1126/science.abg0821 (2021).

11. Dowall, S. D. et al. Elucidating variations in the nucleotide sequence of Ebola virus associated with increasing pathogenicity. Genome Biol 15, 540, doi:10.1186/PREACCEPT-1724277741482641 (2014).

12. Dong, X. et al. Variation around the dominant viral genome sequence contributes to viral load and outcome in patients with Ebola virus disease. Genome Biol 21, 238, doi:10.1186/s13059-020-02148-3 (2020).

13. Salguero, F. J. et al. Comparison of rhesus and cynomolgus macaques as an infection model for COVID-19. Nat Commun 12, 1260, doi:10.1038/s41467-021-21389-9 (2021).

14. Moore, S. C. et al. Amplicon-Based Detection and Sequencing of SARS-CoV-2 in Nasopharyngeal Swabs from Patients With COVID-19 and Identification of Deletions in the Viral Genome That Encode Proteins Involved in Interferon Antagonism. Viruses 12, doi:10.3390/v12101164 (2020).

15. Nasir, J. A. et al. A Comparison of Whole Genome Sequencing of SARS-CoV-2 Using Amplicon-Based Sequencing, Random Hexamers, and Bait Capture. Viruses 12, doi:10.3390/v12080895 (2020).

16. Caly, L. et al. Isolation and rapid sharing of the 2019 novel coronavirus (SARS-CoV-2) from the first patient diagnosed with COVID-19 in Australia. Med J Aust 212, 459–462, doi:10.5694/mja2.50569 (2020).

17. Prince, T. et al. Sequence analysis of SARS-CoV-2 in nasopharyngeal samples from patients with COVID-19 illustrates population variation and diverse phenotypes, placing the in vitro growth properties of B.1.1.7 and B.1.351 lineage viruses in context. bioRxiv, 2021.2003.2030.437704, doi:10.1101/2021.03.30.437704 (2021).

18. Wang, B. et al. Resistance of SARS-CoV-2 variants to neutralization by convalescent plasma from early COVID-19 outbreak in Singapore. NPJ Vaccines 6, 125, doi:10.1038/s41541-021-00389-2 (2021).

19. Saad-Roy, C. M. et al. Epidemiological and evolutionary considerations of SARS-CoV-2 vaccine dosing regimes. Science 372, 363–370, doi:10.1126/science.abg8663 (2021).

20. Gomez-Carballa, A., Bello, X., Pardo-Seco, J., Martinon-Torres, F. & Salas, A. Mapping genome variation of SARS-CoV-2 worldwide highlights the impact of COVID-19 super-spreaders. Genome Res 30, 1434–1448, doi:10.1101/gr.266221.120 (2020).

21. Diez-Fuertes, F. et al. A Founder Effect Led Early SARS-CoV-2 Transmission in Spain. J Virol 95, doi:10.1128/JVI.01583-20 (2021).

22. Tasakis, R. N. et al. SARS-CoV-2 variant evolution in the United States: High accumulation of viral mutations over time likely through serial Founder Events and mutational bursts. PLoS One 16, e0255169, doi:10.1371/journal.pone.0255169 (2021).

23. Ward, T. et al. Growth, reproduction numbers and factors affecting the spread of SARS-CoV-2 novel variants of concern in the UK from October 2020 to July 2021: a modelling analysis. BMJ Open 11, e056636, doi:10.1136/bmjopen-2021-056636 (2021).

24. Rader, B. et al. Crowding and the shape of COVID-19 epidemics. Nat Med 26, 1829–1834, doi:10.1038/s41591-020-1104-0 (2020).

25. Kraemer, M. U. G. et al. Spatiotemporal invasion dynamics of SARS-CoV-2 lineage B.1.1.7 emergence. Science 373, 889–895, doi:10.1126/science.abj0113 (2021).

26. Korber, B. et al. Tracking Changes in SARS-CoV-2 Spike: Evidence that D614G Increases Infectivity of the COVID-19 Virus. Cell 182, 812–827 e819, doi:10.1016/j.cell.2020.06.043 (2020).

27. Goh, K. C. et al. Molecular determinants of plaque size as an indicator of dengue virus attenuation. Sci Rep 6, 26100, doi:10.1038/srep26100 (2016).

28. Kato, F. et al. Characterization of large and small-plaque variants in the Zika virus clinical isolate ZIKV/Hu/S36/Chiba/2016. Sci Rep 7, 16160, doi:10.1038/s41598-017-16475-2 (2017).

29. Wang, W. et al. SARS-CoV-2 nsp12 attenuates type I interferon production by inhibiting IRF3 nuclear translocation. Cell Mol Immunol 18, 945–953, doi:10.1038/s41423-020-00619-y (2021).

30. Munday, D. C. et al. Interactome analysis of the human respiratory syncytial virus RNA polymerase complex identifies protein chaperones as important cofactors that promote L-protein stability and RNA synthesis. J Virol 89, 917–930, doi:10.1128/JVI.01783-14 (2015).

31. Noton, S. L., Aljabr, W., Hiscox, J. A., Matthews, D. A. & Fearns, R. Factors affecting de novo RNA synthesis and back-priming by the respiratory syncytial virus polymerase. Virology 462-463, 318–327, doi:10.1016/j.virol.2014.05.032 (2014).

32. Martin, M. Cutadapt removes adapter sequences from high-throughput sequencing reads. *EMBnet*. journal 17, 10–12 (2011).

33. Joshi, N. & Fass, J. (2011).

34. Dong, X. et al. Variation around the dominant viral genome sequence contributes to viral load and outcome in patients with Ebola virus disease. Genome biology 21, 1–20 (2020).

35. Kim, D., Langmead, B. & Salzberg, S. L. HISAT: a fast spliced aligner with low memory requirements. Nature methods 12, 357 (2015).

36. Li, H. et al. The sequence alignment/map format and SAMtools. Bioinformatics 25, 2078–2079 (2009).

37. Morelli, M. J. et al. Evolution of foot-and-mouth disease virus intra-sample sequence diversity during serial transmission in bovine hosts. Veterinary research 44, 12 (2013).

38. Dong, X. et al. Identification and quantification of SARS-CoV-2 leader subgenomic mRNA gene junctions in nasopharyngeal samples shows phasic transcription in animal models of COVID-19 and aberrant pattens in humans. bioRxiv (2021).

39. Au, C. H., Ho, D. N., Kwong, A., Chan, T. L. & Ma, E. S. BAMClipper: removing primers from alignments to minimize false-negative mutations in amplicon next-generation sequencing. Scientific reports 7, 1–7 (2017).

40. Buchrieser, J. et al. Syncytia formation by SARS-CoV-2-infected cells. EMBO J 39, e106267, doi:10.15252/embj.2020106267 (2020).

41. Thi Nhu Thao, T. et al. Rapid reconstruction of SARS-CoV-2 using a synthetic genomics platform. Nature 582, 561–565, doi:10.1038/s41586-020-2294-9 (2020).

